# Remodeling of epigenome and transcriptome landscapes with aging in mice reveals widespread induction of inflammatory responses

**DOI:** 10.1101/336172

**Authors:** Bérénice A. Benayoun, Elizabeth A. Pollina, Param Priya Singh, Salah Mahmoudi, Itamar Harel, Kerriann M. Casey, Ben W. Dulken, Anshul Kundaje, Anne Brunet

**Affiliations:** Department of Genetics, Stanford University School of Medicine, Stanford, CA 94305, USA; Current affiliation: Leonard Davis School of Gerontology, University of Southern California, Los Angeles, CA 90089, USA; USC Norris Comprehensive Cancer Center, Los Angeles, CA 90089; USC Stem Cell Initiative, Los Angeles, CA 90089; Harvard Medical School, Boston, MA 02115, USA; Silberman Institute of Life Sciences, The Hebrew University of Jerusalem, Givat Ram, Jerusalem, 91904, Israel; Department of Comparative Medicine, Stanford University School of Medicine, Stanford, CA 94305, USA; Paul F. Glenn Laboratories for the Biology of Aging, Stanford University, Stanford, CA 94305, USA

## Abstract

Aging is accompanied by the functional decline of tissues. However, a systematic study of epigenomic and transcriptomic changes across tissues during aging is missing. Here we generated chromatin maps and transcriptomes from 4 tissues and one cell type from young, middle-age, and old mice, yielding 143 high-quality datasets. We focused specifically on chromatin marks linked to gene expression regulation and cell identity: histone H3 trimethylation at lysine 4 (H3K4me3), a mark enriched at promoters, and histone H3 acetylation at lysine 27 (H3K27ac), a mark enriched at active enhancers. Epigenomic and transcriptomic landscapes could easily distinguish between ages, and machine learning analysis showed that specific epigenomic states could predict transcriptional changes during aging. Analysis of datasets from all tissues identified recurrent age-related chromatin and transcriptional changes in key processes, including the upregulation of immune system response pathways such as the interferon signaling pathway. The upregulation of interferon response pathway with age was accompanied by increased transcription of various endogenous retroviral sequences. Pathways deregulated during mouse aging across tissues, notably innate immune pathways, were also deregulated with aging in other vertebrate species – African turquoise killifish, rat, and humans – indicating common signatures of age across species. To date, our dataset represents the largest multi-tissue epigenomic and transcriptomic dataset for vertebrate aging. This resource identifies chromatin and transcriptional states that are characteristic of youthful tissues, which could be leveraged to restore aspects of youthful functionality to old tissues.

## Introduction

The functional decline of organs and tissues is a hallmark of aging, and it is accompanied by changes in gene expression and chromatin modifications across cell types and tissues (Benayoun et al. 2015; Booth and Brunet 2016; Pal and Tyler 2016; Sen et al. 2016). Aging is the primary risk factor for a variety of chronic diseases, including neurodegeneration, cardiovascular disease, diabetes, osteoporosis, and cancer. Several conserved pathways are deregulated during aging, defining hallmarks or pillars of aging (Lopez-Otin et al. 2013; Kennedy et al. 2014). One such hallmark is the accumulation of epigenetic alterations, defined in this context as changes to gene regulation by chromatin modifications. Perturbation in chromatin modifying enzymes can extend lifespan in invertebrate models (Benayoun et al. 2015; Pal and Tyler 2016; Sen et al. 2016), suggesting that loss of chromatin homeostasis may drive aspects of aging. Chromatin marks are relatively stable and can even persist through cell division (Kouskouti and Talianidis 2005).Therefore, sustained alterations to the chromatin landscape may mediate, at least in part, the propagation of age-associated functional decline.

Changes in chromatin marks (*e.g.* DNA methylation, histone modifications) have been observed throughout life in multiple species and tissues (Benayoun et al. 2015; Booth and Brunet 2016; Pal and Tyler 2016; Sen et al. 2016). However, most of this knowledge has relied on DNA methylation or global assessments of histone modification changes (*e.g.* mass spectrometry, western-blot, or immunostaining) rather than locus-specific evaluation (*e.g.* ChIP-seq) (Horvath 2013; Benayoun et al. 2015; Wagner 2017). Several genome-wide studies have interrogated locus-specific changes in histone modifications, chromatin states, as well as changes in gene expression in several cell and tissue types with mammalian aging (*e.g.* adult stem cells, liver, pancreatic beta-cells, neurons, T cells) (Rodwell et al. 2004; Cheung et al. 2010; Liu et al. 2013; Shulha et al. 2013; Bochkis et al. 2014; Sun et al. 2014; Avrahami et al. 2015; White et al. 2015; Zheng et al. 2015; Moskowitz et al. 2017; Stegeman and Weake 2017; Ucar et al. 2017; Nativio et al. 2018). However, while these studies have provided important insights into genome-wide chromatin and transcriptome remodeling with age, they have remained restricted to specific cell types and/or a mark. Thus, whether general rules and patterns govern age-related chromatin and transcriptional changes – and how they are linked –across tissues, cell types, and organisms remains largely unknown.

Here, we investigate genome-wide changes to transcription and chromatin during aging in mouse, focusing on chromatin marks linked to transcriptional activation and cell identity: H3K4me3, a mark of active/poised promoters (Heintzman et al. 2007), and H3K27ac, a mark of active enhancers (Heintzman et al. 2007). We generated epigenomic and transcriptomic maps of the heart, liver, cerebellum, olfactory bulb and primary cultures of neural stem cells (NSCs) throughout mouse lifespan (youth, middle age, and old age), a resource comprising in total 143 high-quality datasets. Our analysis identified age-related remodeling of both transcriptional and chromatin landscapes across tissues. Using machine-learning models, we identify chromatin-level predictors of transcriptional remodeling with age. We observe a recurrent upregulation of interferon-related signaling pathways at the chromatin and transcriptional levels, which is concomitant with the transcriptional upregulation of transposable elements and endogenous retroviruses with aging. Finally, we test conservation of these age-related changes across vertebrate species. This resource identifies conserved epigenomic and transcriptional signatures during vertebrate aging, the understanding of which will be critical to restore old tissues to a more youthful and healthy state.

## Results

### A genome-wide epigenomic and transcriptomic landscape of in four tissues and one cell type during mouse aging

To understand how chromatin and transcriptional profiles change across multiple tissues during aging, we collected tissues and cells from C57BL/6 male mice at 3 different time points throughout their life: youth [3 months], middle age [12 months] and old age [29 months]. We focused on a subset of tissues (*i.e.* heart, liver, cerebellum, olfactory bulb) that are known to display age-related functional decline (Enwere et al. 2004; Sussman and Anversa 2004; Zhang et al. 2010; Shioi and Inuzuka 2012; Mobley et al. 2014; Delire et al. 2016), and that are clearly anatomically defined. We also derived primary cultures of neural stem and progenitor (NSCs) from these young, middle-aged, and old mice. For each tissue or cell culture from all 3 ages, we generated transcriptomic maps (RNA-seq) and epigenomic maps (ChIP-seq of total Histone 3 distribution [H3] for normalization, trimethylation of Histone 3 at lysine 4 [H3K4me3], and acetylation of Histone 3 at lysine 27 [H3K27ac]), yielding a total of 143 high-quality datasets (**Figure 1A-B, S1A; Supplementary Table S1**). We chose H3K4me3 and H3K27ac because both chromatin marks are associated to ‘active chromatin’ because spread of both marks have been associated with information about cell identity and specific transcriptional states. Indeed, H3K4me3 is preferentially enriched at active promoters and H3K27ac is preferentially enriched at active enhancers (Heintzman et al. 2007). In addition to H3K4me3 intensity (*i.e.* ChIP-seq signal per base pair), broad H3K4me3 domains mark genes that are important for cell identity and function (Bernstein et al. 2006; Benayoun et al. 2014; Chen et al. 2015) and exhibit increased transcriptional levels (Chen et al. 2015) and consistency (Benayoun et al. 2014). In addition to H3K27ac intensity, large clusters of H3K27ac-enriched enhancers, known as Super Enhancers (Hnisz et al. 2013) or Stretch Enhancers (Parker et al. 2013), mark enhancers of genes that are cell-or tissue-specific and highly transcribed in that specific cell or tissue. Importantly, ChIP datasets for the H3K4me3 and H3K27ac histone mark were normalized to paired total Histone H3 ChIP-seq data, to account for potential changes in local nucleosome landscape with age. Whenever possible, tissues from age-matched mice were examined by a trained histopathologist to record age-related changes and account for any irregularities in the samples (**Figure S1B**).

**Figure 1:**
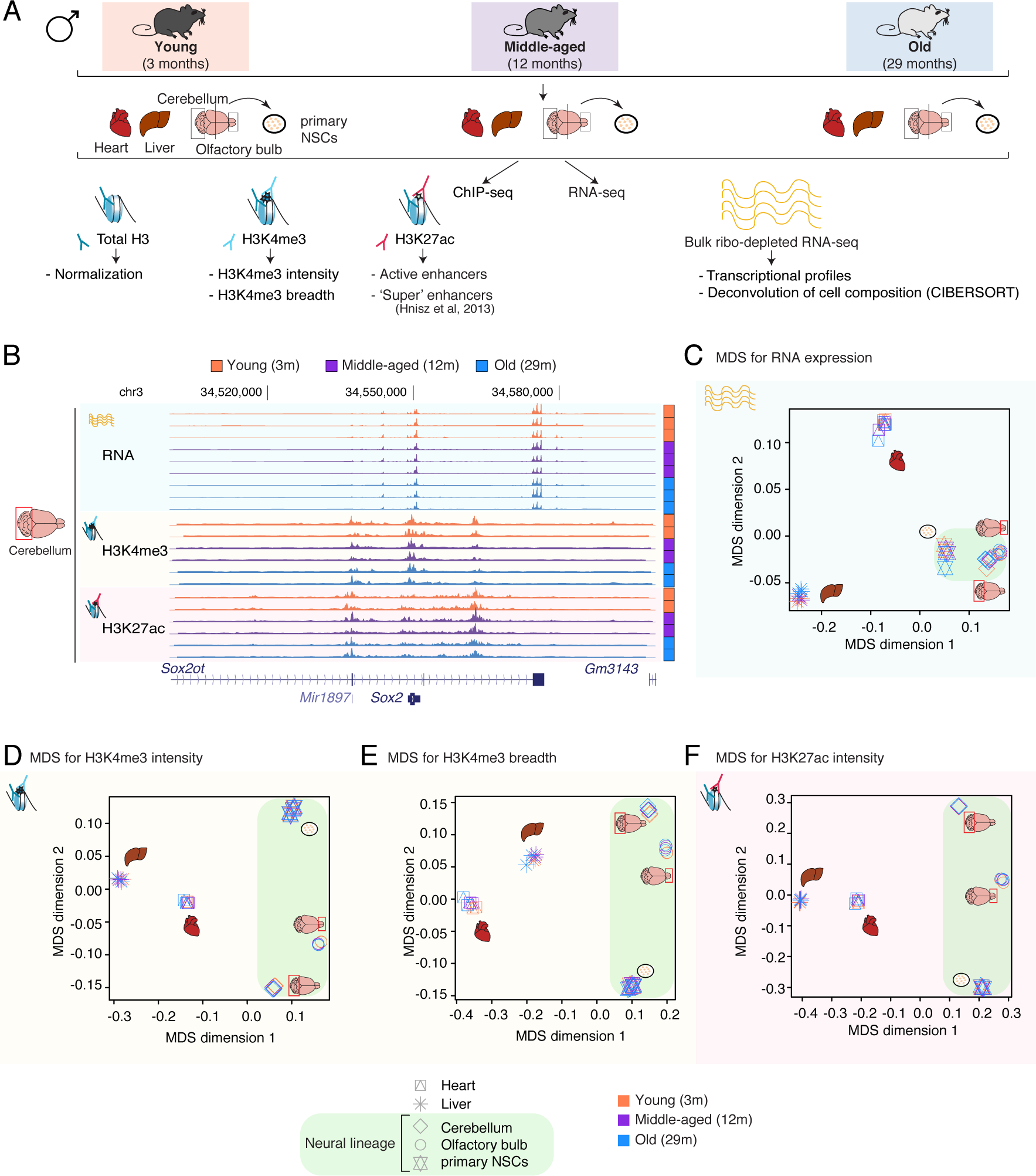
A genome-wide epigenomic and transcriptomic landscape of in four tissues and one cell type during mouse aging. (A) Experimental data setup. Also see Supplementary Table S1. (B) Example UCSC genome browser region showing tracks of generated datasets in cerebellum tissue. (C-F) Multidimensional Scaling analysis results across datasets based on RNA expression (C), H3K4me3 peak intensity (D), H3K4me3 peak breadth (E), or H3K27ac peak intensity at all peaks (F).

To visualize the degrees of similarity of our genomic samples, we used Multidimensional scaling (MDS) (Chen and Meltzer 2005). MDS allows the projection of high-dimensionality ‘omics’ data on a set number of dimensions to facilitate visualization of data similarities (Chen and Meltzer 2005). MDS analysis on RNA, H3K4me3 intensity, H3K4me3 breadth, H3K27ac intensity, or H3K27ac breadth revealed that, as expected, the main source of sample separation corresponds to the nature of the tissue, regardless of the age of the animal (**Figure 1E-F, S1D-E**). Principle component analysis (PCA), another dimensionality reduction method (Ringner 2008), yielded very similar results when extracting the first 2 components (**Figure S1F-G**). Thus, we used MDS analysis for all subsequent analyses. Our results are consistent with the fact that RNA profiles, H3K4me3 and H3K27ac are associated with cell identity (Hnisz et al. 2013; Benayoun et al. 2014; Wagner et al. 2016), and indicate that overall tissue and cell identities remain quite stable during aging.

To understand how age impacts the global epigenomic and transcriptomic landscapes in each tissue or cell type, we performed MDS one tissue/cell type at a time (**Figure 2A-J, S2A-O**). Interestingly, in all tissues and for all features (RNA, H3K4me3 intensity and breadth, H3K27ac intensity and breadth), there was a clear progressive separation based on the age of the samples of origin, with the young samples clustering closer to the middle-age samples, and further away from the old samples (**Figure 2A-J, S2A-O**). For primary NSC cultures, there was also a clear separation with age for H3K4me3 intensity, H3K4me3 breadth, and H3K27ac Super Enhancers (**Figure S2F, S2I,** and **S2O**). However, the transcriptome and H3K27ac intensity of NSCs (**Figure S2C** and **S2L**) did not separate well with respect to age, possibly because of technical noise. Together, these results indicate that genome-wide RNA and features of H3K4me3 and H3K27ac deposition can distinguish between ages in different tissues and cells.

**Figure 2:**
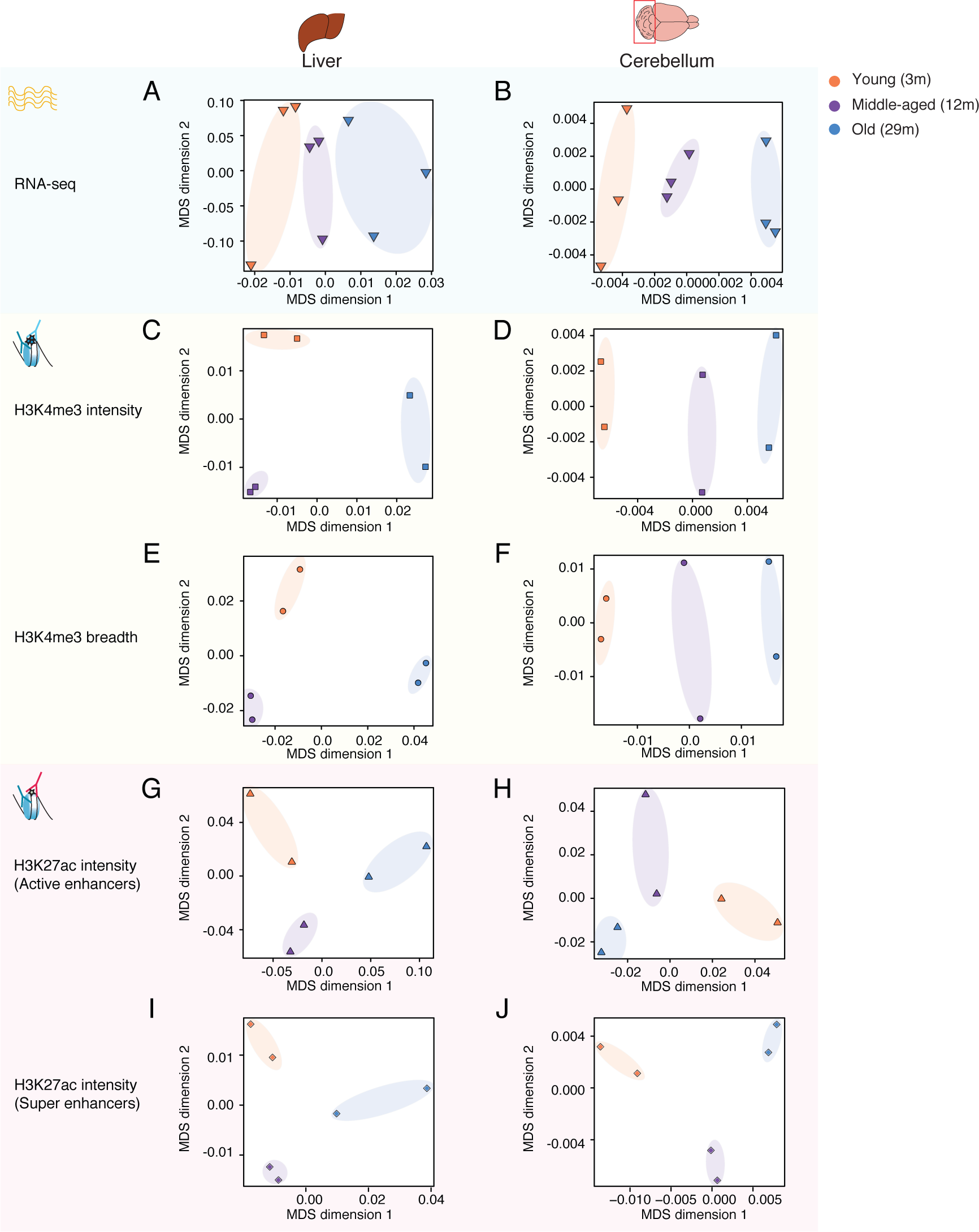
Separation of samples across tissues and cell types as a function of age. Multidimensional Scaling analysis results across samples derived from specific tissues, Liver and Cerebellum, based on RNA expression (A-B), H3K4me3 peak intensity (C-D), H3K4me3 peak breadth (E-F), H3K27ac peak intensity (all peaks: G-H; Super Enhancers only: I-J).

### Machine-learning reveals that age-related epigenomic changes can predict transcriptional changes

To understand how age-related changes in the epigenome predict age-related transcriptional changes, we leveraged the power of machine-learning (**Figures 3, S3**). Using four types of algorithms (*i.e.* neural networks [NNET], support vector machines [SVM], gradient boosting [GBM] and random forests [RF]), we trained machine learning models to discriminate between transcriptional changes with age (upregulated, downregulated, or unchanged gene expression during aging) (**Figure 3A**). As potential predictors for the models, we used, for each gene, specific features from chromatin datasets generated for this study (*e.g.* amount of H3K4me3 signal at the promoter, change of H3K4me3 breadth awith age; see methods), from mouse ENCODE ChIP-seq datasets in heart, liver, cerebellum and olfactory bulb in young mice (*e.g.* Pol2, CTCF, H3K27me3, H3K4me1) (Shen et al. 2012; Yue et al. 2014) (**Supplementary Table S2**), and from the underlying DNA sequence (*e.g.* %CpG in promoter, exon number, *etc.*) [See methods for details on all used features]. Because absolute gene expression levels could influence the ability to call differential gene expression (Oshlack and Wakefield 2009), we also included, as a potential predictor, the average expression level of the genes in the young samples, expressed in FPKM (fragments per read per million).

**Figure 3:**
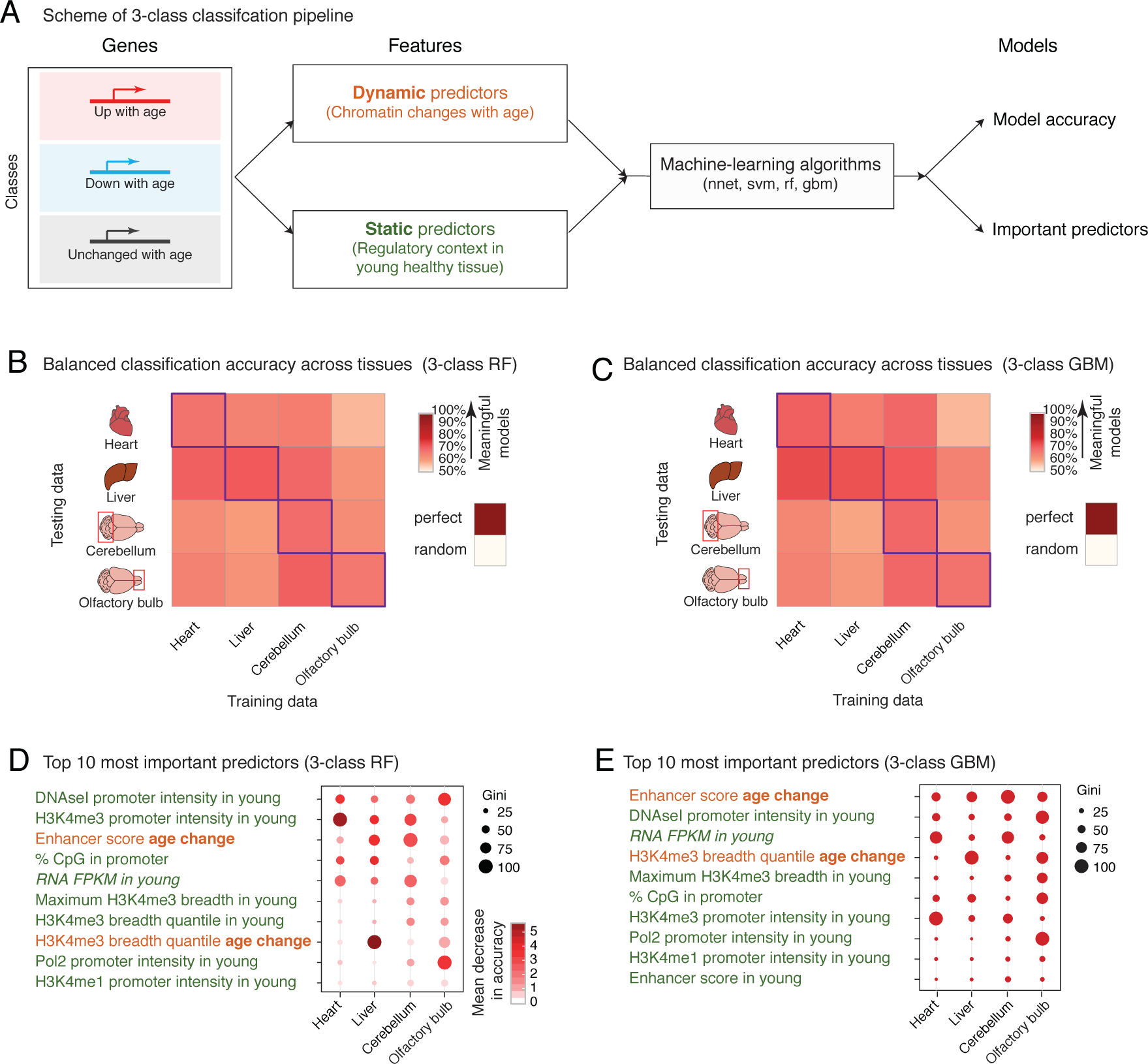
Machine-learning analysis reveals that changes in enhancer score and H3K4me3 domain breadth with age can predict transcriptional aging. (A) Scheme of the 3-class machine learning pipeline. Nnet: neural network, svm: support vector machine, rf: random forest, gbm: gradient boosting machine. (B-C) Balanced classification accuracy over the 3-classes (*i.e.* upregulated, downregulated, and unchanged genes) cross tissues for Random Forest models (B) or Gradient Boosting models (C). The accuracy of the model trained in a specific tissue on the same tissue (*e.g.* the liver-trained model on liver data) is measured using held-out validation data, and for cross-tissue validation, the entire data of the tested tissue was used. ‘Random’ accuracy is displayed to illustrate the accuracy of a meaningless model (∼50%). All tests were significantly more accurate than random. The robustness of the prediction is supported by the fact that samples for RNA and chromatin profiling were collected from independent mice at 2 independent times (**Supplementary Table S1**). Balanced accuracy across the 3-classes is reported. (D-E) Feature importance from Random Forest models (D; Gini score and mean decrease in accuracy) or Gradient Boosting models (E; Gini score). High values indicate important predictors. Also see analysis of 2-class models (*i.e.* upregulated *vs.* downregulated genes) in **Figure S3**. Note that 2-class models, though containing less biological information, outperformed 3-class models, which is consistent with the increased complexity of a classification problem with the number of classes to discriminate.

All four machine learning models assigned genes to the correct transcriptional change with age (*e.g.* upregulation, etc.) 66-79% of the time on average, significantly above that of a random classification (50%) (**Supplementary Table S3, Figure 3B-C**). Models derived using tree-based algorithms (*i.e.* GBM and RF) performed slightly better than the other models (69-79% *vs.* 66-76%) (**Supplementary Table S3**). The accuracy was similar whether validation was performed within or across tissues (**Supplementary Table S3, Figure 3B-C**). These results support the idea that epigenomic remodeling is associated with transcriptional remodeling during aging. These observations also suggest that genes that are dysregulated with age share common epigenomic signatures even if they are found at different loci in different tissues. Interestingly, important predictors of age-related transcriptional changes in all tissues were dynamic changes in enhancer score (*i.e.* change in the amount of H3K27ac detected at enhancers during aging) and dynamic changes in the breadth of the associated H3K4me3 domains with aging (**Figure 3D-E, S3D-E**). Other predictors of the transcriptional status of genes during aging were static, describing the young chromatin context (*e.g.* promoter accessibility by DNAseI, H3K4me3 promoter intensity or H3K4me3 domain breadth) (**Figure 3D-E**). To note, we cannot fully exclude that this finding may result from incomplete accounting for gene expression levels differences, as H3K4me3 and promoter accessibility have been associated to higher expression levels (Consortium 2012). Nevertheless, this observation raises the intriguing possibility that the chromatin context of a gene in a youthful context might predict at least in part change in expression of that gene with age, perhaps because active loci are impacted by accumulated stresses and injuries throughout life.

Together, these data indicate that chromatin states can predict age-dependent changes in transcription.

### Immune response pathways are robustly upregulated during mouse aging

We next asked which genes or pathways were significantly deregulated with age across tissues (**Figure 4, S4**). At the transcriptional level, we identified only 16 upregulated genes (and 0 downregulated) with aging in all tissues, when each tissue was assessed separately (FDR < 5%; **Figure 4A**). These genes encode complement and coagulation factors (*i.e.* C1qa, C1qc, C4) and several interferon-response related proteins (*i.e.* GBP6, GBP10, IFI27l1, IFI44, IFIT3, IFITM3), as well as a protein known to play a key role in leukocyte transendothelial migration (*i.e.* ITGB2) (Guan et al. 2015). Though the observed overlap is small, these results suggest that a common response to aging across tissues could be linked to an immune response.

**Figure 4:**
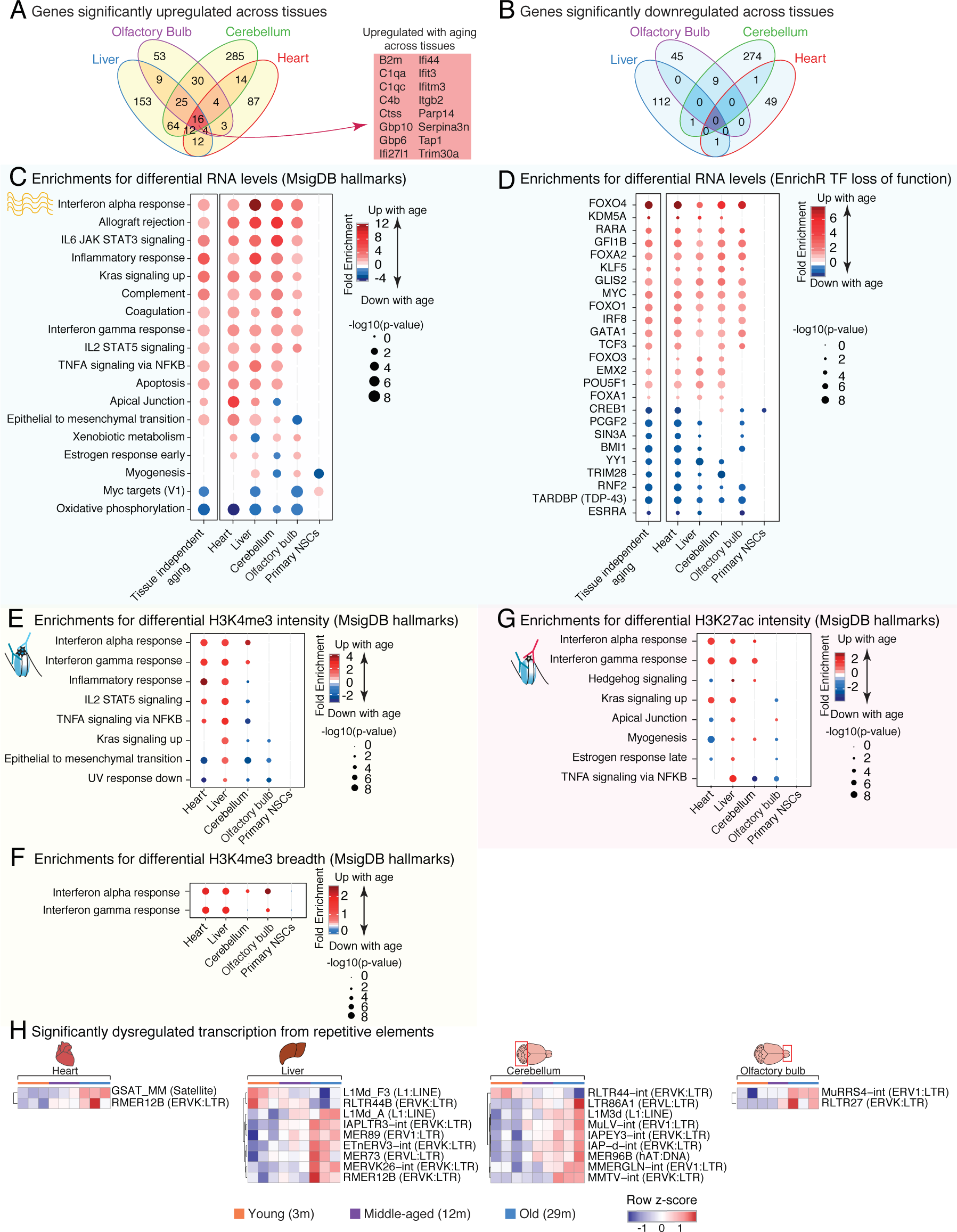
Age-misregulated pathways reveal the activation of an innate immune signature. (A-B) Venn diagram for the overlap of significantly upregulated (A) or downregulated (B) genes with aging in each tissue called by DESeq2 at FDR < 5%. (C-F) Functional enrichments using the minimum hypergeometric [mHG] test at for differential RNA expression (C-D), for differential H3K4me3 intensity (E), for H3K4me3 breadth (F) and for differential H3K27ac intensity (G). Enriched pathways were plotted if 4 out of the 6 tests (RNA) or 3 out of the 5 tests (chromatin marks) were significant (FDR < 5%). (H) Heatmap of expression for repetitive elements with significant differential expression with aging (TE-transcript quantification and DESeq2 statistical test at FDR < 5%).

To explore this hypothesis further, we leveraged the statistical power of the whole dataset and performed an analysis of age-related transcriptional changes using all tissues and ages combined, but including tissue of origin as a covariate. This analysis identified 945 genes showing tissue-independent age-related change, with 771 genes upregulated and 174 genes downregulated with age at FDR < 5% (**Figure S4F; Supplementary Table S4**). Consistently, among these 771 upregulated genes were 14 of the 16 genes commonly upregulated using the analysis in each tissue (see **Figure 4A**). Since it enables a more general view of our RNA-seq data, we include this global analysis in addition to tissue-specific analysis for functional enrichment analyses below (labeled “tissue-independent aging”).

For H3K4me3 and H3K27ac chromatin marks, we did not find any recurrently misregulated epigenomic loci across tissues at FDR < 5%, which is compatible with the tissue-specific nature of regulatory elements, where the same genes can have different enhancers or alternative promoters in different tissues where they are expressed. However, it cannot be excluded that this difference could result from different sensitivity of the read-out to changes in chromatin vs. tramscriptome measurements, including because of potential changes in cell composition (see below).

Next, we investigated whether we could identify recurrently misregulated pathways and gene sets with age across tissues (**Figure 4C-G, S4G-I**). Rank statistics analysis identified several pathways that were recurrently deregulated with aging, not only at the transcriptome level, but also at the level of chromatin modifications (**Figure 4C, 4E-G, S4G-I**). Down-regulated pathways included pathways related to mitochondrial dysfunction (*e.g.* ‘oxidative phosphorylation’, ‘TCA cycle’), compatible with previous work using RNA microarrays in mouse and human tissues (Zahn et al. 2006; Zahn et al. 2007). Upregulated pathways included protein homeostasis (*e.g.* ‘lysosome’, ‘ribosome’) or immune signaling pathways (*e.g.* ‘Inflammatory response’, ‘Interferon alpha response’, ‘Interferon gamma response’, ‘Chemokine signaling pathway’) (**Figure 4C, 4E-G, S4G-I**), which is consistent with previous observations in a number of aging tissues (Stegeman and Weake 2017), such as choroid plexus tissue (Baruch et al. 2014) or kidney (Rodwell et al. 2004; O’Brown et al. 2015). Complement and coagulation-related pathways were also significantly upregulated across tissues and cell types (**Figures 4C, S4G**). However, the strongest signal came from the interferon alpha and gamma response pathways, which were significantly induced across aged tissues at the transcriptional and chromatin levels (**Figure 4C, 4E-G, S4G-I**). Notably, we confirmed the transcriptional activation of the interferon response by Ingenuity Pathway Analysis (e.g. IFNG, IFNB1, IFNAR; **Supplementary Table S5A**). While this age-related inflammatory response has been observed at the transcriptional level across many studies (Stegeman and Weake 2017), this is the first time this is observed at the transcriptional and chromatin levels.

Interferon response activation can stem from (i) a response to exogenous viral infection, (ii) reactivation of endogenous transposable elements (TEs) with aging (De Cecco et al. 2013; Wood and Helfand 2013), and/or (iii) more generally, aberrant cytosolic DNA detection by the Cyclic GMP-AMP synthase pathway (*i.e.* cGAS) (Sun et al. 2013; West et al. 2015). Since both old and young mice were kept in SPF facilities at Charles River and at Stanford, and were documented to not have viral infection upon routine testing, we queried TE expression in the different tissues during aging using our RNA-seq datasets. Increased activity of TEs has been reported with aging in several species (*i.e.* worm, mouse, humans) and cell types (Maxwell et al. 2011; De Cecco et al. 2013; Wood and Helfand 2013; Van Meter et al. 2014). Using the ‘TE-transcript’ and HOMER repeats pipelines, we identified repetitive elements whose transcription levels are significantly changed with aging (**Figure 4H; Supplementary Table S6A-E**). Consistently with previous reports, the majority of elements with changed transcriptional levels were upregulated during aging (**Figure 4H**). Moreover, most significantly upregulated elements belonged to <u>e</u>ndogenous retro<u>v</u>irus (ERV) families (**Figure 4H**).

The interferon signaling pathway upregulation is also compatible with the significant upregulation of the KEGG 2017 ‘cytosolic DNA-sensing pathway’ genes, which corresponds to cGAS activation (**Figure S4C**). The cGAS pathway has been found to be upregulated in senescent cells due to aberrant cytoplasmic chromatin (Dou et al. 2017) and in response to deficient mitochondrial DNA – a known consequence of aging (West et al. 2015). Thus, activation of the cGAS pathway by endogenous DNA may play a role in the age-related increase in the interferon response.

Consistent with functional pathway enrichment results, targets genes of pro-inflammatory transcription factors IRF8 and TCF3 were significantly upregulated (**Figure 4D**). Consistently, targets of pro-inflammatory transcription factors IRF3, IRF5 and IRF7 were signitifantly induced across tissues according to Ingenuity Pathway Analysis (**Supplementary Table S5A**). FOXO targets were also significantly upregulated with aging (**Figure 4D**). As FOXO factors are known to be pro-longevity genes (Martins et al. 2016) and to modulate innate immunity (Seiler et al. 2013), this upregulation may result from a compensatory mechanism and could contribute to the upregulation of the innate immune response with aging. In addition, Myc targets were also upregulated (**Figure 4D**), consistent with Myc’s reported pro-aging effects (Hofmann et al. 2015). Finally, targets of the RNA binding protein TARDBP (also known as TDP-43) were significantly downregulated with aging across tissues (**Figure 4D**). Mutations in human *TARDBP/TDP-43* are involved in the pathogenesis of amyotrophic lateral sclerosis (ALS) and frontotemporal dementia (FTD) (Scotter et al. 2015), and TDP-43 has been suggested to play a role in retrovirus suppression by host cells (Ou et al. 1995) and in microglia activation (Zhao et al. 2015). Together, these results suggesting that dysregulation of targets of several transcription factors and RNA binding proteins could also be a key player in the upregulation of innate immune response pathways with aging.

Finally, we asked if the transcriptional increase in immune pathways in tissues over the course of aging could result from the transcriptome of infiltrated immune cells (Rodwell et al. 2004; Lumeng et al. 2011; Pinto et al. 2014; O’Brown et al. 2015). Using CIBERSORT to perform computational deconvolution of aging tissue RNA-seq datasets (Newman et al. 2015), no significant change could be detected in the proportions of predicted inflammatory cell signatures (**Figure S5** and **Supplementary Table S7**). However, expression levels of known immune markers revealed were slightly upregulated with age (**Figure S5F-G**). Thus, at least a portion of the observed inflammatory response with age might be due to a low, but increased, amount of infiltrated immune cells in old tissues.

### Conservation of mouse age-regulated transcriptional trends across vertebrate model organisms

To investigate whether the age-related changes observed across mouse tissues were conserved in other vertebrate species, we used publicly available aging transcriptome datasets in other species, including rat (Yu et al. 2014), human (Mele et al. 2015), and the naturally short-lived African turquoise killifish (Baumgart et al. 2014; Baumgart et al. 2016). We identified rat, human and turquoise killifish orthologs for each mouse gene that was significantly deregulated with aging. Notably, the interferon alpha and gamma response pathways were also significantly misregulated during aging in rat, human, and turquoise killifish samples (**Figure 5A, S6A**). In addition, we also determined the transcriptional trajectory with aging of genes orthologous to the genes changes with aging in mouse tissues (**Figure 5B, S6B**). In general, similar aging trajectories (*i.e.* upregulation with age or downregulation with age) were observed for the same genes in similar tissues across vertebrate species (**Figure 5B, S6B**). These trajectories were overall less strongly conserved in the GTEx human data, perhaps because other factors (*e.g.* environmental differences, presence of specific diseases) may overshadow aging differences in human tissues. Indeed, when accounting for body mass index and metabolic disease-status in an independent human liver microarray dataset (GSE61260) (Horvath et al. 2014), gene expression trajectories with aging were more similar between mouse and human (**Figure S6C-D**). Collectively, these data indicate that core signatures of innate immune responses are consistently upregulated with aging across species.

**Figure 5:**
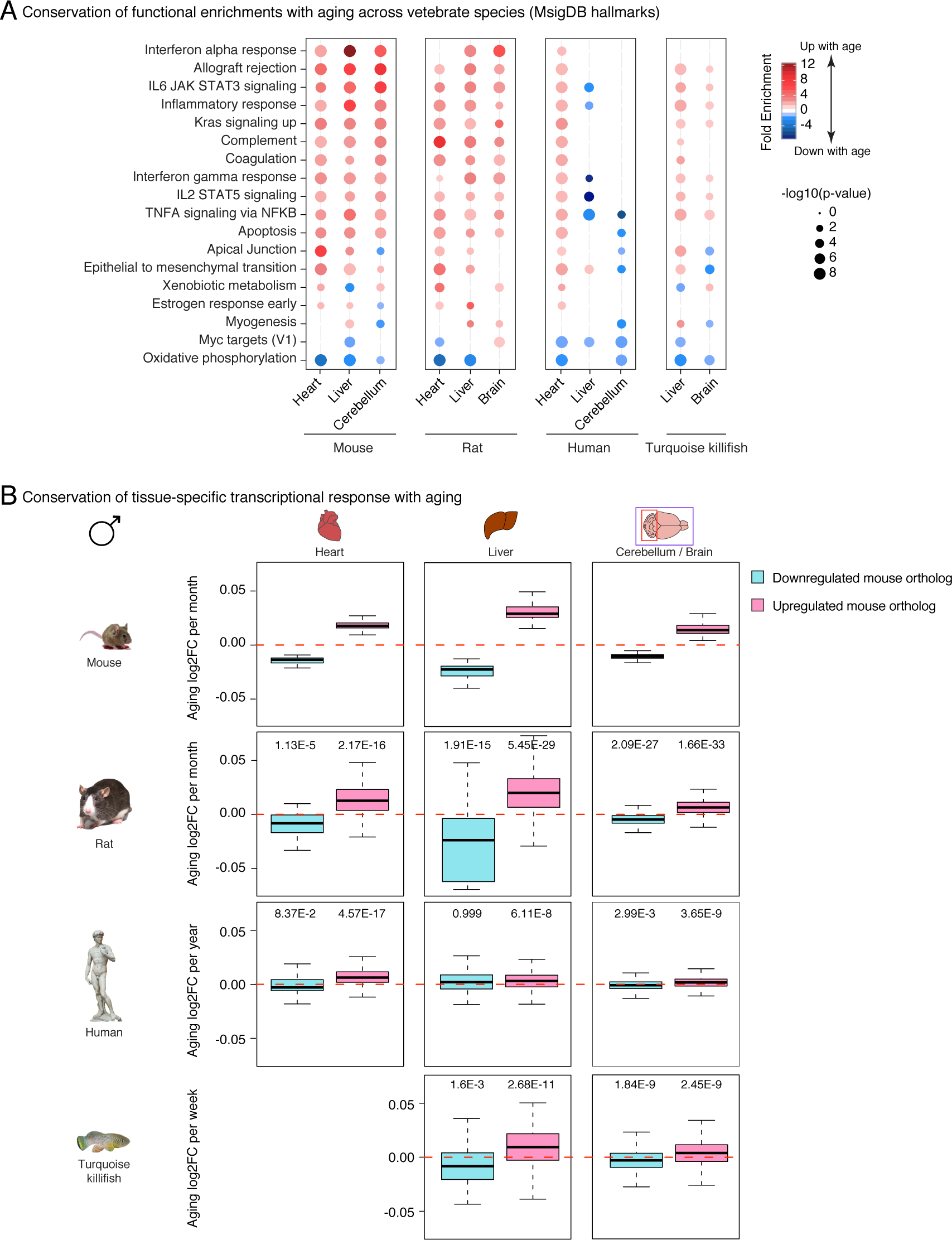
Age-related transcriptional deregulation from mouse are overall conserved across vertebrate species. (A) Functional enrichments using the mHG test at for differential RNA expression with aging in mouse, rat, human and African turquoise killifish samples. The mouse data is a subset of the panel shown in **Figure 4C** and is plotted as a comparison point. (B) Log2 Fold Change per unit of time during aging for genes orthologous to differentially expressed mouse genes in this study in rat, human and African turquoise killifish samples [see methods]. The mouse data is plotted for comparison. Significance in on sample one-sided Wilcoxon test the fold changes and 0 (*i.e.* no change with age). Only data from males is plotted here. Data with the contribution of female samples (when available) is in **Figure S6A**.

## Discussion

To understand the effect of aging on genomic regulation and chromatin identity with aging, we have generated transcriptomic and epigenomic maps in young, middle-aged, and old mice from a variety of tissues and cells known to show functional decline with aging (*i.e.* heart, liver, cerebellum, olfactory bulb and cultured primary NSCs). To our knowledge, this dataset represents the largest epigenomic and transcriptomic dataset for mammalian aging to date and will serve as a resource for the aging community. Thanks to the inclusion of a middle-age time point, we find that progressive changes accumulate throughout mouse lifespan not only at the transcriptional level, but also at the level of several chromatin features. The progressive accumulation of remodeling of histone modifications with aging is reminiscent of the DNA methylation clock paradigm (Horvath 2013; Cole et al. 2017; Quach et al. 2017; Stubbs et al. 2017; Wang et al. 2017). Thus, the existence of these progressive changes is compatible with the existence of chromatin modification clocks. Additional time points and individuals will be required to build such clocks and compare their performance to that of the well-established DNA methylation clock. The potential functional interaction between these different levels of molecular clocks could provide important insights on the regulation of cellular and organismal aging.

### Machine learning as a powerful tool to study aging epigenomics

By leveraging the power of machine-learning, we show that observed age-related epigenomic remodeling is predictive of age-related transcriptional changes. This is consistent with the ‘histone code hypothesis’, whereby the chromatin context may direct transcriptional outputs (Jenuwein and Allis 2001), and supports the idea that the rules that govern the relationship between the chromatin landscape and transcriptional outputs are generally preserved throughout life. To note, these models cannot be used to provide clues about the flow of information between chromatin and transcriptional changes. Thus, we cannot exclude that age-related transcriptional changes may precede observed chromatin changes. However, our models do indicate that the changes at the chromatin and transcription are to some degree coordinated, and that the breakdown of gene regulation levels is a complex occurrence. Important identified predictors of the relationship between transcriptional and chromatin aging may present interesting entry point for future mechanistic studies of aging epigenomics.

### Innate immune pathways are broadly induced during aging across tissues and species

Our analyses reveal that immune pathways are upregulated with aging across tissues and species. This increase in immune activity in the absence of exogenous pathogens is consistent with the importance of the concept of inflamm-aging (Xia et al. 2016). Notably, we find that the interferon response pathways, both alpha and gamma, are recurrently and robustly activated with aging across vertebrate tissues. Although interferon signaling is traditionally associated with the response to viral infection, the interferon pathway can also be induced in response mitochondrial DNA stress and cytosolic DNA detection (Sun et al. 2013; West et al. 2015), including the reactivation of endogenous transposable elements (De Cecco et al. 2013; Wood and Helfand 2013). Accordingly, our analyses find evidence for induction of cytosolic DNA-sensing pathway genes, as well as a significant transcriptional upregulation of several families of endogenous transposable elements (TEs). Many TEs can retain key viral characteristics, including the ability to replicate, to form viral particles and to potentially trigger host immune responses (Kassiotis and Stoye 2016). Thus, our results are compatible with the intriguing possibility that the global increase in innate immunity signals across tissues during aging might be mediated, at least partly, through the detection of endogenous aberrant DNA or reactivated endogenous retroviral particles. In addition, the impact of infiltrated immune cells will need further studies, in particular at single cell level, in order to disentangle the relative contribution of infiltrated immune cells and endogenous detection of aberrant DNA.

### A resource for the study of aging genomics

To date, this dataset is one of the largest existing aging multi-omics datasets and could be integrated to future studies with additional marks. It is one of the rare cases with a middle-aged point, in addition to a young and old time-points, which helps understand epigenomic and transcriptomic aging as trajectories rather than end-point results. The transcriptomic arm of our dataset is consistent with the wealth of other published transcriptional aging datasets using microarrays and RNA-seq technologies (Stegeman and Weake 2017), with an increase in inflammation, stress responses and a decrease in mitochondria function. Several studies have started to interrogate genome-wide chromatin remodeling with vertebrate aging (Cheung et al. 2010; Liu et al. 2013; Shulha et al. 2013; Bochkis et al. 2014; Sun et al. 2014; Avrahami et al. 2015; Zheng et al. 2015; Moskowitz et al. 2017; Ucar et al. 2017), with concomitant changes in the transcriptional landscape. However, the combination of cross-tissue assessment (*i.e.* heart, liver, cerebellum, olfactory bulb and cultured primary NSCs), multiple chromatin feature profiling (*i.e.* total H3, H3K4me3 and H3K27ac) and though youth, middle and old age is unique, and enabled us to conduct an integrated study of conserved and coordinated genomic misregulation with aging. To note, our datasets have primarily focused on histone modifications associated with active/accessible chromatin. However, other features of chromatin, including heterochromatin, are also likely to play important roles in epigenome remodeling during aging and could bring important insights. Finally, we have generated these profiles only in males thus far. Future work will need to assess the impact of sex-dimorphism, as well as diverse genetic background, in the regulation of transcriptional and chromatin aging. Further mining of this dataset by the community should help identify candidate regulators that affect age-dependent dysfunction across multiple tissues in vertebrates.

## Methods

### Mouse husbandry

All animals were treated and housed in accordance to the Guide for Care and Use of Laboratory Animals. All experimental procedures were approved by Stanford’s Administrative Panel on Laboratory Animal Care (APLAC) and were in accordance with institutional and national guidelines. Male C57BL/6 mice at different ages (3, 12 and 29 months) were obtained from the National Institute on Aging (NIA) colony at Charles Rivers, which is an SPF facility, and were acclimated at the SPF animal facility at Stanford University for 1-2 week before processing. All animals were euthanized between 10-12am for tissue harvesting or NSCs isolation. No animals were censored.

### Tissue dissection and histopathology

For our aging ‘omics’ studies, we selected the heart, liver, cerebellum, olfactory bulb, as well as primary NSCs cultures derived from the Subventricular Zone (SVZ) because (i) these tissues and cells are known to display age-related functional decline (Enwere et al. 2004; Sussman and Anversa 2004; Zhang et al. 2010; Shioi and Inuzuka 2012; Mobley et al. 2014; Delire et al. 2016) and (ii) the tissues are all clearly defined anatomically, which guarantees reproducible isolation across animals of different ages and minimizes the risk that observed differences comes from dissection differences. For the heart, we dissected the ventricle tissue for chromatin and RNA extraction. For the liver, the top-most lobe was harvested for chromatin and RNA extraction. The complete anatomical structure of the cerebellum and olfactory bulb (both sides) were used for chromatin and RNA extraction. Detailed information for each sample are reported in **Supplementary Table S1**.

Heart and liver tissue were harvested for histopathology evaluation in parallel to RNA-seq profiling. Tissues samples were processed as described previously (Harel et al. 2015). Briefly, tissues were collected into ice cold PBS, washed, then fixed for 24h in Bouin’s solution at room temperature. The following day, tissues were rinsed in PBS and paraffin-embedded using standard procedures. Sections of 5 µm were stained with hematoxylin and eosin (HE) and Mason’s trichrome for histopathologic analysis. All tissues were reviewed by a board-certified veterinary anatomic pathologist who was blinded to animal identification (K.C.). Microscopically, cross-sectional areas of the liver and heart were qualitatively evaluated for age-related histopathologic lesions from 3 months old (n=3), 12 months old (n=3) and 29 months old (n=3) C57BL/6 male mice. Liver tissues were specifically evaluated for hepatic capsule contour, perivascular lymphoid aggregates, hepatocellular atrophy, extramedullary hematopoiesis (foci of myeloid and erythroid precursor cells), anisocytosis (variation in hepatocellular size), anisokaryosis (variation in hepatic nuclear size), bile duct proliferation, fibrosis, sinusoidal cellularity, and the presence of Ito cells (*i.e.* major resident cell type involved in liver fibrosis, also known as hepatic stellate cells). Because of dissection of parts of the tissue for RNA-seq analysis, representative histologic sections of all heart chambers (*i.e.* left and right atria, left and right ventricles), valves, and vessels could not be consistently obtained for all mice. Thus, comparisons could not be drawn between specific heart chambers. Based on tissue availability, heart samples were evaluated for age-related pathology including myocardial fibrosis, myocardial degeneration (*i.e.* loss of cross striations, vacuolization, necrosis), myocardial inflammation, lymphoid aggregates, mineralization, and atrial thrombosis.

Livers of 3 months old mice were characterized by smooth hepatic capsules and rare scattered foci of extramedullary hematopoiesis (**Figure S1B**). Normal lobar architecture and sinusoidal organization of the liver were evident, with evenly spaced centrilobular and portal regions. At 12 months of age, in addition to rare foci of extramedullary hematopoiesis, small perivascular lymphoid aggregates were identified in the liver. Slight anisocytosis and anisokaryosis were also observed across hepatocytes at 12 months of age. By 29 months of age, foci of extramedullary hematopoiesis and perivascular lymphoid aggregates were increased in number and size in the liver. Undulation of the hepatic capsule was evident and was considered a direct result of hepatic cord atrophy. Anisocytosis and anisokaryosis were more pronounced in 29 month-old mice than in 12 months old mice. Additional histologic findings that were present only in the livers from 29 months old mice included occasional bile duct hyperplasia, increased sinusoidal cellularity (notably for Kupffer cells), increased numbers of lipid-rich Ito cells, and rare sinusoidal fibrosis (**Figure S1B**). Overall, no observable histologic differences were identified between available heart samples from 3 months, 12 months, and 29 months old mice. Occasionally, small foci of myocardial degeneration (characterized by loss of myofiber cross-striation and vacuolization) were noted across all age groups. Surprisingly, hemangiosarcoma (an endothelial cell neoplasm) was noted within the left ventricle of one mouse within the 29-month old cohort. The tumor comprised ill-defined and infiltrative vascular channels lined by variably plump neoplastic endothelial cells. Lymphatic dilation and perivascular edema were noted adjacent to the neoplasm in the heart and were thus considered sequelae to tumor formation, rather than an age-related manifestation. The corresponding RNA-seq sample did not encompass the tumor, which is compatible with the absence of gross outlier behavior in the RNA-seq analysis. Since the whole cerebella and olfactory bulb samples were used to extract RNA, the histological status of these tissues could not be assessed in our cohorts.

### Primary NSC cell culture

NSCs were isolated from 3 months, 12 months, and 29 months old male C57BL/6 mice from the NIA aging colony at Charles River as previously described (Renault et al. 2009; Webb et al. 2013). Briefly, the subventricular zone (SVZ) was finely microdissected and chopped into ice-cold PBS. Microdissected SVZs from 5 age-matched mice were pooled from each age group to make a single culture. Tissue chunks were digested by 10 min incubation 14 U/mL Papain (Worthington) at 37^°^C. Following mechanical trituration, cells were purified by a 22% Percoll (GE Healthcare) gradient. NSCs were plated at a density of <10^5^ cells/cm^2^ as non-adherent spheres in NSC growth media (*i.e.* Neurobasal A medium [Life technologies] medium supplemented with 1% penicillin/streptomycin/glutamine [Life technologies], 2% B27 supplement [Life technologies] and 20 ng/mL each of FGF2 [Peprotec] and EGF [Peprotec]). Cultures from all ages were plated at the same density to minimize differences in autocrine/paracrine signaling. Cells were passaged using Accutase enzyme (Stem Cell Technologies, 07920). All datasets were generated on cells grown as non-adherent spheres that had been disassociated the day prior, replated as a suspension culture and collected at the end of passages 2-3 (14 days of culture).

NSCs cultures derived from 3m, 12m and 29m old animals were assessed at the end of passages 1 and 3 (**Figure S1C**). This analysis revealed that there were significantly fewer NSCs in 29m cultures compared to 3m cultures at passage 1, although this difference seemed to have been erased by passage 3 (**Figure S1C**).

### Tissue isolation and chromatin preparation

ChIP experiments on different tissues were performed as follows: olfactory bulbs were microdissected from 3 months, 12 months, and 29 months old males C57BL/6 mice (NIA colony at Charles Rivers) and weighed (**Supplementary Table S1**). Olfactory bulbs were pooled from 4-8 mice of the same age per biological replicate. Cerebella were dissected, weighed, and pooled from 2 mice of the same age per biological replicate. The entire upper left lobe of the liver was dissected and weighed from an individual mouse and was used for a single biological replicate. Following removal of blood and aorta, the heart ventricles from an individual mouse were dissected and weighed from an individual mouse and served as a single biological replicate. All tissue samples were finely minced with a fresh razor blade, then resuspended in ice-cold PBS. Following mincing, tissues were crosslinked via addition of 1% formaldehyde for 15min at room temperature and quenched by the addition of 0.125M glycine for 5min at room temperature. ChIP experiments on mouse primary NSC cultures were performed as previously described (Benayoun et al. 2014). Briefly, NSCs neurospheres (passages 2-3) (**Supplementary Table S1**) were dissociated 16-18 hours prior to collection. Cells were crosslinked at a density of 100,000 cells per mL in ice cold PBS with 1% formaldehyde for 9min at room temperature, and the crosslinking reaction was quenched with 0.125M glycine for 5min at room temperature.

For tissues and cells, after the quenching step, all crosslinked samples were washed three times with 1X PBS containing protease inhibitors cocktail (Roche). Samples were then snap frozen in liquid nitrogen, and preserved at −80^°^C as dry cell or tissue pellets until the day of the IP. On the day of the IP, samples were resuspended in 1mL of SDS lysis buffer (50 mM Tris-Hcl pH7.5, 10 mM EDTA, 1% SDS in PBS at pH7.4) containing protease inhibitors cocktail (Roche). Chromatin was sheared with a Vibra-Cell Sonicator VC130 (Sonics) 8 times [tissues] or 7 times [cells] for 30 seconds at 60% amplitude with probe model CV188, and then diluted 1:5 fold in RIPA buffer (1% NP-40, 0.5% Sodium deoxycholate in PBS, pH 7.4) containing protease inhibitors cocktail (Roche).

### Chromatin quantification and immunoprecipitation

For liver, heart, and cerebellum, chromatin content was measured and equalized for all ages to enable fair comparison across samples of a tissue. To measure the chromatin content of sonicated samples, samples were incubated with 0.2 μg/mL RnaseA (Life Technologies) for 1h at 37^°^C and 0.2 mg/mL Proteinase K (Life Technologies) for for 1h at 55^°^C, and crosslinks were reversed by incubation at 80^°^;C for 2 hours. DNA was precipitated by the addition of 0.1 volume 3M sodium acetate, 2μg glycoblue (Life Technologies), and 2.5 volumes of 100% ethanol. The resulting DNA concentration was quantified by Nanodrop technology.

We used 20μg of chromatin for the H3 ChIPs, 50μg for the H3K4me3 ChIPs, and respectively 70μg (heart) or 100μg (liver and cerebellum) for the H3K27ac ChIPs. For the olfactory bulb, chromatin from approximately 30mg of tissue was used for immunoprecipitation with anti-H3 antibody, and 60mg was used for immunoprecipitation with anti H3K4me3 and H3K27ac antibodies. For adult NSCs, we used chromatin from ∼250,000 cells for the H3 ChIP, ∼750,000 cells for the H3K4me3 ChIP, and ∼1,000,000 for the H3K27ac ChIP. ChIP was performed as previously described (Webb et al. 2013; Benayoun et al. 2014). The corresponding amount of chromatin diluted in RIPA buffer containing protease inhibitors cocktail (Roche) [see above] was incubated overnight at 4^°^C with the following antibodies: 5μg H3K4me3 antibody (Active Motif #39159, lot #1609004), 5μg Histone H3 (Abcam #1791 lot #GR178101-1), and 7μg H3K27ac (Active motif #39133, lot #1613007).

### Next-generation sequencing ChIP library generation

For olfactory bulb libraries and the first set of NSCs libraries, libraries were generated with the Illumina Tru-Seq kit (#IP-202-1012) according to the manufacturer instructions. Briefly, repaired and adapter ligated DNA was size selected in range of 250-400bp and PCR amplified for 16 (H3), 17 (H3K4me3) and 18-19 (H3K27ac) cycles. Because the Illumina Tru-Seq kit became backordered during the course of this study, we generated libraries using the NEBNext DNA library prep kit (E6040L) for the liver, heart, cerebellum, and the second set of H3 and H3K4me3 NSCs libraries, according to the manufacturer instructions. Repaired and adapter ligated DNA was size selected in range of 250-400bp using agarose gel electrophoresis and PCR amplified for 14 (H3), 16-17 (H3K4me3) or 17-18 (H3K27ac) cycles. Library quality was assessed using the Agilent 2100 Bioanalyzer (Agilent Technologies). Single-end 101bp reads were generated on Illumina HiSeq2000 machines, and subsequently analyzed with our standardized ChIP-seq data analysis pipeline [see below].

### S2 cell culture for ChIP normalization control

We explored the use of spike-in for ChIP normalization in some of our samples. Indeed, ChIP-seq from spiked-in chromatin from a different species (e.g. *Drosophila*) (termed ChIP-Rx for ChIP with reference exogenous genome (ChIP-Rx) has been shown to allow genome-wide quantitative comparisons of histone modification status across cell populations (Orlando et al. 2014). For example, the fraction of histone modification ChIP-seq reads (*e.g.* H3K4me3) corresponding to the species of interest (*e.g.* mouse) compared to the exogenous control (*e.g. Drosophila*) can be used to detect global differences in the amount of that histone modification between conditions of interest (*e.g.* aging). As several studies suggested that global levels of total histone proteins or specific histone modifications may change with aging (Feser et al. 2010; O’Sullivan et al. 2010; Liu et al. 2013), we thought that ChIP-Rx could be helpful to account for this potential change.

To this end, D*rosophila* S2 cells were grown at 25°C in Schneider’s media (Invitrogen) with 10% heat inactivated fetal bovine serum and 1% penicillin/streptomycin/glutamine (Invitrogen). Cells were crosslinked at a density of 1-2×10^6^ cells/ml in 1X PBS with 1% formaldehyde for 8min at room temperature, and the reaction was quenched with 0.125M glycine for 5min at room temperature. Next, 10^7^ S2 cells were sonicated 7 times for 30s at 60% amplitude in 1mL of SDS lysis buffer (50 mM Tris-Hcl pH7.5, 10 mM EDTA, 1% SDS in PBS pH7.4) containing protease inhibitors cocktail (Roche) using a Vibra-Cell Sonicator VC130. The S2 chromatin was added to heart, liver, and cerebellum tissues at a ratio of 1:120μg of chromatin. For the second set of NSCs ChIPs, S2 chromatin was added at a ratio of 1:4 cell ratio. ChIP was performed using the combination of probed tissue chromatin and S2 chromatin (Orlando et al. 2014). The first set of NSCs ChIPs and the Olfactory bulb ChIPs were performed, before the authors became aware of the possibility to perform ChIP-rx.

However, we found that too few reads per spiked sample mapped to the *Drosophila* genome dm3 in our libraries (<50,000 on average), which may explain why there was more variation n observed percentages of *Drosophila* reads between samples of the same age than between ages. For this reason, we did not end up using the S2 chromatin mappings to normalize our data in the processing pipeline. However, these reads are present in the ChIP and could be used for re-analysis.

### ChIP-Seq data processing

For ChIP-seq data processing, 101 bp reads were trimmed using Trimgalore v0.3.1 (www.bioinformatics.babraham.ac.uk/projects/trim_galore/) to retain high-quality bases with phred score of greater than 15. The trimming command was: Trim_galore -q 15 --stringency 3 --length 36 --phred33 data_file.fastq. Reads were mapped to the mm9 mouse genome assembly using bowtie version 0.12.7 (Langmead et al. 2009). PCR duplicates were removed using FIXSeq (fixseq-e2cc14f19764) (Hashimoto et al. 2014), which accounts for overdispersed per-base read count distributions using a nonparametric method which was shown to substantially improve the performance and precision of ChIP-seq analysis compared with existing alternatives (Hashimoto et al. 2014). Regions of significant enrichment were determined using MACS2 v2.0.8 (Zhang et al. 2008) using the --broad option to enable wider regions of enrichment to be detected. The ‘--keep-dup=all’ option was used to supersede MACS2 basic duplicate removal method since the FIXSeq method had already been applied. Total H3 ChIP-seq samples were used to determine the local background of H3 modification ChIP-seq datasets. Significant ChIP peaks of interest were annotated to the gene with the closest transcription start site in the mm9 assembly using the HOMER suite (Heinz et al. 2010).

### H3K4me3 breadth remodeling analysis

To compare changes in the breadth of H3K4me3 domains, we improved upon our previously developed pipeline to computationally adjust samples such that that the signal to noise ratio across all peaks is equalized between samples (Benayoun et al. 2014). We first created a reference peakset for all comparative analyses using pooled QC reads from all ages and replicates, and MACS2 (v2.08) as a peak calling software as highlighted above (hereafter referred to as ‘metapeaks’). To match the signal-to-noise ratios across all aging samples, we then down-sampled reads separately in each H3K4me3 ChIP-seq biological sample to match the coverage histogram across all samples over the metapeaks intervals, similar to (Benayoun et al. 2014). This procedure matches the “height” of the peaks from the peak caller’s point of view. The appropriate down-sampling rate that allows the coverage histogram of higher sensitivity H3K4me3 ChIP-seq samples to be equal or lower than that of the lowest sensitivity H3K4me3 ChIP-seq sample was determined by minimizing the p-value of Kolmogorov-Smirnov test (comparison to the sample with lowest H3K4me3 ChIP-seq sensitivity). In addition, to limit the effect of variations in input sample depth, we also matched the effective depth of H3 input samples to that of the lowest available sample. Final H3K4me3 domain breadth calls per samples were performed by using MACS2.08 with the same parameters as above. IntersectBed (BedTools-2.16.1) (Quinlan and Hall 2010) was used to estimate the length coverage of the sample peaks over the reference metapeaks. The goal of this pipeline is to increase the likelyhood that called gains/losses of breadth result from a change in breadth of the enriched region, and not simply from an underlying difference in H3K4me3 intensity.

Differential breadth was then estimated using the ‘DESeq2’ R package (DESeq2 1.6.3) (Love et al. 2014).

### Dimensionality reduction techniques for data exploration

To perform Multidimensional Scaling (MDS) analysis, we used a distance based on spearman rank correlation value (1-rho) between samples, which was then provided to the core R command ‘cmdscale’. Principal component analysis (PCA) was performed using the core R command ‘prcomp’. Dimensionality reduction techniques were applied to log2 transformed ‘DESeq2’ VST normalized values.

### H3K4me3 and H3K27ac intensity remodeling analysis

Similar to above, we created a reference peak sets for all comparative analyses using pooled QC reads from all ages and replicates, and MACS2 (v2.08) as a peak calling software as highlighted above (hereafter referred to as ‘metapeaks’). Intensity signals for histone H3 modifications normalized to the local H3 occupancy were obtained using the ‘Diffbind’ R package (Diffbind v1.12.3) (Ross-Innes et al. 2012). Normalized intensities were then used to estimate differential intensities as a function of age using the ‘DESeq2’ R package (DESeq2 v1.6.3) (Love et al. 2014).

### Differential nucleosome calling using total H3 ChIP-seq

To compare changes in the local H3 deposition landscape, we used DANPOS v2.2 (Chen et al. 2013) and DiNUP v1.3 (Fu et al. 2012) nucleosome-calling softwares. For higher confidence results, we considered nucleosomes to be significantly remodeled if the position were called differential by DANPOS (p < 1×10^-15^) and DiNUP (FDR < 0.05) following the same direction (*i.e.* increased *vs.* decreased signal). Consistently with a previously published MNase study on mouse aging liver tissue (Bochkis et al. 2014), we found that significant nucleosome remodeling with chronological aging seems to be restricted to a limited number of loci. Based on our observations and previously published reports, it is possible that decreased nucleosome occupancy may only be a cell-type or context specific effect of aging. Differential nucleosome calls were used as features in our machine learning models [see below].

### Cell and tissue isolation for RNA isolation

For RNA isolation, we used a new cohort of aging male C57BL/6 mice (same ages than the ChIP-seq cohort), and RNA-seq datasets were generated at a later time than the ChIP-seq datasets (see **Supplementary Table S1**).

For RNA extraction on tissues: olfactory bulbs were microdissected from 3 months, 12 months, and 29 months old males C57BL/6 mice and weighed, and tissues from 2 independent mice of the same age were pooled per biological replicate. Cerebellum samples were similarly dissected, weighed, and samples from 2 mice of the same age were pooled per biological replicate. For the liver, the leftmost part of upper left lobe of the liver was dissected and weighed from an individual mouse and was used for a single biological replicate. For the heart, following removal of blood, the bottom-most part of heart ventricles from an individual mouse were dissected, weighed and used as a single biological replicate. All tissue samples were flash-frozen in liquid nitrogen until further processing. Tissues were resuspended in 600μL of RLT buffer (RNAeasy plus mini kit, Qiagen) supplemented with 2-mercaptoethanol as recommended by manufacturer, then homogenized on Lysing Matrix D 2mL tubes (MP Biomedicals) on a FastPrep-24 machine (MP Biomedicals) with a speed setting of 6. Heart tissue was homogenized for 4 times 30 seconds, and all other tissues were homogenized for 40 seconds. Subsequent RNA extraction was performed using the RNAeasy plus mini kit (Qiagen) following the manufacturer’s instructions.

Primary NSC neurospheres (passages 2-3) were dissociated 16-18 hours prior to collection and seeded in 12-well plates. Cells were spun down and collected in RLT buffer (RNAeasy plus mini kit, Qiagen) supplemented with 2-mercaptoethanol as recommended by manufacturer.

### RNA-seq library preparation

1μg of total RNA was combined to 2μL of a 1:100 dilution of ERCC RNA Spike-in mix (Thermo-Fisher scientific) in nuclease-free water, as recommended by the manufacturer. The resulting mix was then subjected to rRNA depletion using the NEBNext rRNA Depletion Kit (New England Bioloabs), according to the manufacturer’s protocol. Strand specific RNA-seq libraries were then constructed using the SMARTer^®^ Stranded RNA-Seq Kit (Clontech # 634836), according to the manufacturer’s protocol. Paired-end 75bp reads were generated on the Illumina NetxSeq 500 platform, and subsequently analyzed with a standardized RNA-seq data analysis pipeline (described below).

### RNA-seq analysis pipeline

Paired-end 75bp reads were trimmed using Trimgalore v0.3.1 (www.bioinformatics.babraham.ac.uk/projects/trim_galore/) to retain high-quality bases with phred score of greater than 15 and a remaining length > 35bp. Read pairs were then mapped to the UCSC mm9 genome build using STAR v2.4.0j (Dobin et al. 2013). Read counts were assigned to genes using the featureCounts functionality of ‘subread’ (subread-1.4.5-p1) (Liao et al. 2014). Read counts were imported into R to estimate differential gene expression as a function of age using the ‘DESeq2’ R package (DESeq2 1.6.3) (Love et al. 2014).

To map repetitive element expression, we used the TE-transcript 1.5.1 software (Jin et al. 2015), with mm9 repeat masker data downloaded from the UCSC table browser. Read counts were imported into R to estimate differential gene expression as a function of age using the ‘DESeq2’ R package (DESeq2 1.6.3) (Love et al. 2014). We also used the ‘analyzeRepeats.pl’ functionality of the HOMER software (Heinz et al. 2010). In that specific case, read counts were imported into R to estimate differential gene expression as a function of age using the ‘DESeq2’ R package (DESeq2 1.16.1) (Love et al. 2014).

### Machine-Learning analysis

Machine-learning models were built for each tissue. No models were built in the NSCs since no gene was found to be significantly dysregulated by RNA-seq. We built classification models in each tissue independently using 4 different classification algorithms as implemented through R package ‘caret’ (caret v.6.0-47). Classification algorithms for neural nets (NNET; ‘pcaNNet’) are directly implemented in the ‘caret’ package. Auxiliary R packages were used with ‘caret’ to implement random forests (RF; ‘randomForest’ v.4.6-12), gradient boosting (GBM; ‘gbm’ v.2.1.1) and radial support vector machines (SVM; kernlab v.0.9-25). ‘Caret’ was allowed to optimize final model parameters on the training data using 10-fold cross validation. Accuracies, sensitivities and specificities for all classifiers in their cognate tissue were estimated using a test set of randomly held out 1/3 of the data obtained using the ‘createDataPartition’ function in ‘caret’ package (**Supplementary Table S3**). Feature importance estimation was only done using random forests and gradient-boosted trees since other algorithms do not natively allow for it. The Gini score for feature importance was computed by ‘caret’ for each feature in the GBM and RF models.

For each expressed gene in each tissue, we extracted two types of ‘features’: (i) **dynamic** features, which encode changes to the chromatin landscape with age, and (ii) **static** features, which reflect the status of the chromatin and transcriptional landscape in young healthy animals. For dynamic features, we included: number of H3 nucleosomes with increased occupancy between 3 and 29m, number of H3 nucleosomes with decreased occupancy between 3 and 29m, maximum log2 fold change in H3 occupancy between 3 and 29m, slope of H3K4me3 intensity change at annotated promoter with aging, slope of H3K27ac intensity change at annotated promoter with aging, slope of H3K4me3 breadth quantile change at annotated promoter with aging, and slope of H3K27ac intensity at stitched enhancers with aging (*i.e.* Enhancer score).

For static features using the data we generated, we included the enhancer presence and type (*i.e.* None, Typical or Super), the distance of the closest Super Enhancer to the transcriptional start site of the gene, the broad H3K4me3 domain status (*i.e.* None, Typical or Broad), the H3K4me3 breadth quantile in the young sample, the breadth of the broadest H3K4me3 domain associated to gene in the young sample, the average promoter intensity for H3K4me3 and H3K27ac in the young samples (promoters defined as -300bp;+300bp with regards to the transcriptional start sites defined in mm9 build, downloaded from the UCSC genome browser on 2016/1/21), the Enhancer score in the young sample (as defined in (Hnisz et al. 2013)). In addition, we took advantage of the mouse ENCODE datasets (Shen et al. 2012; Yue et al. 2014) in the same tissues (heart, liver, cerebellum and olfactory bulb) in 2 months old mice to select additional potentially informative features (see complete list of datasets **Supplementary Table S2A**). Using the H3K4me1 and H3K27me3 ENCODE ChIP-seq datasets, we collected the average promoter intensity for these marks (same promoter definition as above). In addition, we intersected our young H3K4me3 ChIP-seq datasets to the H3K27me3 peaks to identify whether a gene had a potentially bivalent domain (*i.e.* with overlapping H3K27me3 and H3K4me3 peaks) (Bernstein et al. 2006) in the young animals or not. Using the Pol2 ChIP-seq datasets, we included the number of Pol2 peaks associated with each gene, the absolute distance of the closest Pol2 peak to the transcriptional start site of the gene, the maximal MACS2 score for Pol2 associated to the gene, the average promoter intensity for Pol2. We also extracted the Traveling Ratio (TR) of Pol2, which gives a measure of Pol2 pausing, as described before (Benayoun et al. 2014). Using the CTCF ChIP-seq datasets, we included the number of CTCF peaks associated with each gene, the absolute distance of the closest CTCF peak to the transcriptional start site of the gene, the maximal MACS2 score for CTCF associated to the gene, and the average promoter intensity for CTCF. We also included the average promoter intensity for DNAseI signals, which quantifies how accessible the gene promoter is to the transcriptional machinery. Finally, we included several DNA sequence features from the mm9 build associated to each gene: the presence of a CpG island, the CG and CpG percentage in promoters computed using HOMER, as well as the number of exons in each gene.

Because absolute gene expression levels can influence the ability of differential gene expression pipelines to identify differentially expressed genes (Tarazona et al. 2011), we also included the average expression level of the genes <u>in the young samples</u>, expressed in FPKM (fragments per read per million). This should allow us to take into account potential non-biological effects (*i.e.* effects due to differential expression pipelines) in our models. Any additional feature that is significantly predictive of differential gene expression with age should then be associated independently of the expression level effect, since the expression level is taken into account by the model at the same time when evaluating the importance of features.

We then trained machine-learning classification models in two different ways: (i) either only comparing genes called by DESeq2 as up *vs.* down with age at FDR < 10% (<u>2-class models</u>) or (ii) comparing genes called by DESeq2 as up, down, or unchanged with age at FDR < 10% (<u>3-class models</u>) (**Figure 3 and S3**). These 2 types models allow us to ask different questions: the former interrogates whether are there chromatin differences between upregulated and downregulated genes, without taking the rest of the genes into account, whereas the latter also attempts to identify differences between the genes that change and the ones that do not change during aging. To note, 2-class models systematically outperformed 3-class models in terms of model accuracy (**Figures 3B,C and S3B,C; Supplementary Table S3**), which is consistent with the increased complexity of a classification problem with the number of classes to discriminate.

The relative number of genes that change transcriptionally with age (< 500) is low compared to the number of expressed genes (>3000) (class sizes in each tissues can be found in **Supplementary Table S3**). Thus, to avoid biases in the 3-class classification output due to the imbalanced number of genes called as transcriptionally changed and unchanged with aging, we generated a list of unchanged genes for each sampling that are equal in number to the changed genes in the training set of the cognate tissue (*e.g.* sample 500 unchanged genes to be compared to 500 changed genes in each iteration). We repeated the sampling procedure 100-250 times to eliminate random sampling biases (NNET and SVM: 100 samplings; RF, GBM: 250 samplings). Notably, to improve the signal-to-noise ratio and decrease the inclusion of false negatives (*i.e.* genes called as “unchanged” because of lack of statistical power) in our dataset in the learning and testing of the machine-learning, the constant class was restricted to genes with a log2-fold change (as called by DEseq2) within half a standard deviation of the log2-fold change distribution both above and below 0 (no change).

### Functional enrichment analysis using the minimum hypergeometric test

To perform functional enrichment analysis, we leveraged the minimum HyperGeometric (mHG) distribution, which has been spearheaded through the GOrilla enrichment tool (Eden et al. 2007; Eden et al. 2009). The mHG tests is performed using a ranked list of genes, and allows computing of an exact p-value for the observed enrichment, taking threshold multiple testing into account without the need for simulations, which enables rigorous and rapid statistical analysis of thousands of genes and thousands of functional enrichment terms (Eden et al. 2007; Eden et al. 2009). We used the R implementation of the mHG distribution (‘mHG’ v1.1 package) to run a minimum-hypergeometric (mHG) test as described in (Eden et al. 2007) on gene sets from MSigDB hallmarks (Liberzon et al. 2015), KEGG 2017 (Kanehisa et al. 2017), obtained using KEGG API, after excluding disease pathways so as to focus on core biological processes (see KEGG pathways list with and without disease pathways removed in **Supplementary Table S5B,C**), and transcription factor loss-of-function targets compiled by EnrichR (Kuleshov et al. 2016). The MSigDB hallmarks are highly curated datsets that allow for clean identification of modified pathways, KEGG is an independent curated list of biological pathways, and the EnrichR loss-of-function targets is an unbiased reository derived from published loss-of-function experiments allows to identify unbiasedly potential upstream regulators of observed transcriptional changes with age.

For analysis of function enrichments associations on chromatin datasets, we ranked domains to run the mHG tests based on the intensity or breadth of the mark of interest, and annotated each domain to the gene with the closest transcriptional start site. Genes were often associated to more than one chromatin domain, and only the most extreme domain (*i.e.* more intense, or broadest) were retained when running the mHG enrichment test.

### Ingenuity Pathway Analysis

Upstream regulator analysis was performed using QIAGEN’s Ingenuity Pathway analysis (IPA QIAGEN Redwood City) software, using the genes that passed the filter in our datasets as reference genome, the cutoff of 0.05 for FDR-corrected p-values and species parameter was restricted to mouse. Results in **Supplementary Table S5** are shown for gene sets that passed significance in IPA in at least 4 of the analyzed sets (each tissues and tissue-independent analysis).

### Tissue RNA-seq deconvolution using CIBERSORT

Raw data from RNA-seq datasets from purified mouse cell types was downloaded from the SRA repository. All samples were processed using a standardized pipeline, which consisted of mapping with STAR v2.4.0j (Dobin et al. 2013) to the mm9 assembly, and summarization of reads over mm9 genes using the featureCounts functionality of the subread software suite (subread-1.4.5-p1) (Liao et al. 2014). Read counts from all samples were imported into R for further processing and normalization.

First, single-cell RNA-seq datasets were aggregated into pseudo-bulks by adding reads coming from 4-150 cells of the same cell type from the same study (depending on sequencing depth and number of available cells). Next, any bulk or pseudo bulk sample with more than 18,000 genes without any read detected were eliminated from further processing as too low coverage. Then, all retained quality-checked samples were normalized using the DESeq2 variance stabilizing transformation (VST), and subjected to log2 transformation before upload to the CIBERSORT portal (Newman et al. 2015). The CIBERSORT website was used with default parameters to build the signature matrix and to analyze RNA-seq samples from cell mixtures.

To perform the deconvolution process, CIBERSORT requires an input matrix of reference gene expression signatures (or ‘signature matrix’), collectively used to estimate the relative proportions of each cell type of interest. To provide this reference framework for CIBERSORT, we curated >1500 RNA-seq datasets of purified mouse cell types (**Figure S5A; Supplementary Table S2B**). After undergoing standardized processing, these datasets were used to build a signature matrix for CIBERSORT (**Supplementary Table S7A**). To test the accuracy of the trained signature matrix, we randomly withheld one RNA-seq sample per cell type with at least 3 biological samples for validation. We verified that CIBERSORT was generally sensitive enough to detect various cell types on their own, and within *in silico* mixes of cell types expected to co-occur in tissues (**Figure S5B-C; Supplementary Table S7B**). Using CIBERSORT and the trained signature matrix, we could detect a macrophage/microglia signature in the brain tissue of Alzheimer’s mouse models with known increased inflammatory cell content (Gjoneska et al. 2015; Iaccarino et al. 2016) (**Figure S5D; Supplementary Table S7C**). However, when applied to our RNA-seq datasets, CIBERSORT did not identify significant change in the presence of inflammatory cell signatures, and no strong trend could be observed (**Figure S5E; Supplementary Table S7D**).

### Rat and turquoise killifish RNA-seq processing

Tissue aging RNA-seq datasets from female and male rats were obtained from the Rat Bodymap project (GEO accession number GSE53960) (Yu et al. 2014). Reads were mapped using Kallisto (version 0.43.0) (Bray et al. 2016). DEseq2 normalized fold-changes were then used to estimate differential gene expression as a function of age using the ‘DESeq2’ R package (DESeq2 1.6.3). Orthology tables between rat and mouse genes for aging trends comparison were obtained from ENSEMBL Biomart (2017/05/15).

The African turquoise killifish (*Nothobranchius furzeri,* wild-derived MZM-0410 strain) aging RNA-seq dataset was obtained from GEO (accession number: GSE69122) (Baumgart et al. 2014; Baumgart et al. 2016). Reads were mapped to African turquoise killifish reference transcriptome (GRZ) from NCBI Annotation Release 100 (https://www.ncbi.nlm.nih.gov/genome/annotation_euk/Nothobranchius_furzeri/100) using kallisto. Read pseudo-counts for each transcript were merged and read counts per gene were obtained using Bioconductor R package ‘tximportData’ (Love M, 2017, version 1.6.0). Linear modeling of the differential expression analysis from 5-weeks to 39-weeks old tissues was performed using the R ‘DESeq2’ package as described above. To note, biological replicate clustering by hierarchical clustering and PCA was not very clear, suggesting potential unknown covariates in the dataset that we could not model in our analyses. High confidence orthologs between African turquoise killifish and mouse genes were obtained using bidirectional best BLASTp analysis using longest protein sequences for each of the two species (E-value < 10^-3^).

### Human RNA-seq analysis with aging (GTEx)

Read counts of human tissue RNA-seq with aging per genes were obtained from from the GTEx consortium (v6p version, downloaded 2016/11/22) (Mele et al. 2015). Genotyping principal components, sex, age, as well as other provided sample meta-data (i.e. “RIN” [RNA integrity score], “Ischemic time”, “Fixation time”, “RNA batch”), were also obtained at the time for differential expression analysis. To allow for integration of the discretized age variable into quantitative models, age ranges were converted to a single age at the middle point of the coded range (*e.g.* “20-29” coded as 25 years, “50-59” coded as 55 years). DEseq2 normalized fold-changes were then used to estimate differential gene expression as a function of age (DESeq2 1.6.3). The GTEx v6p count data on genes is reported according to Gencode 19 models, which correspond to Ensembl 74/75 builds (NCBI assembly GRCh37.p13). Using the Biomart mirror for the Ensembl 75 build, we obtained the orthology table between human and mouse genes for aging trend comparisons. We also obtained an independent human liver aging transcriptome dataset (accession GSE61260; “GSE61260_datLiverNormalizedExpr.csv”) (Horvath et al. 2014), along with accompanying sample meta-data (sex, age, BMI, disease status). R package ‘limma’ normalized fold-changes were then used to estimate differential gene expression as a function of age (limma 3.32.10).

Code will be available on GitHub repository.

## Data Access

All new raw sequencing data generated in this study have been deposited at SRA, accession SRP057387 (BioProject PRJNA281127). All accession numbers of previously published datasets can be found in **Supplementary Table S2**.

## Acknowledgements

We thank Aaron Newman and Ash Alizadeh for advice and feedback on the use of CIBERSORT in aging tissue RNA-seq deconvolution and Art Owen for advice and feedback on statistical analysis approaches. We thank Ashley Webb for assistance in the dissection of adult mouse tissues for ChIP-seq and RNA-seq experiments. We thank Katja Hebestreit for helpful advice on functional analyses of the ChIP-seq and RNA-seq experiments. We thank Lauren Booth, Kévin Contrepois, Aaron Daugherty, Changhan D. Lee, Dena Leeman, John Tower, Marc Vermulst, and Robin Yeo for helpful feedback on analyses and manuscript. We thank Matthew Buckley, Brittany Demmitt, Andrew McKay, and Robin Yeo for assisting with independent code-checking.

Illumina HiSeq2000 sequencing was performed at the Stanford Genome Sequencing Service Center, and Illumina NextSeq500 sequencing was performed at the Stanford Functional Genomics Facility. This work was supported in part by NIH P30 CA124435 through the utilization of the following Stanford Cancer Institute Shared Resource, the Genetics Bioinformatics Service Center. This work was also supported by DP1AG044848 (A.B.), P01AG036695 (A.B.), a generous gift from Tim and Michele Barakett (A.B.), and R00AG049934 (B.A.B.).

## Author contributions

B.A.B., E.A.P. and A.B. designed the study. B.A.B. and E.A.P. generated the ChIP-seq and RNA-seq datasets for this study. B.A.B. processed the data and performed the analyses. P.P.S. mapped and quantified the turquoise killifish RNA-seq datasets, generated homology tables between turquoise killifish and mouse genes, and performed independent code-checking. I.H. and B.W.D. helped with dataset generation, and B.W.D. performed independent code-checking. I.H. processed tissues for histological analysis. S.M. performed the Ingenuity Pathway Analysis and performed independent code-checking. K.M.C. performed the histopathology analysis on aging mouse tissues. A.K. advised on data processing pipelines and on the machine learning analyses. B.A.B and A.B. wrote the paper. All authors edited and commented on the manuscript.

## Supplementary Figure legends

**Figure S1: Mouse aging multi-omics dataset across 4 tissues and 1 cell type.**

(A) Summary of the datasets generated in this study. Also see **Supplementary Table S1**. (B) Liver histology of 3 months (i, iv), 12 months (ii, v), and 29 months old (iii, vi) C57BL/6 mice paired to the RNA-seq cohort from this study. Tissue sections were stained with Hematoxylin and eosin, and magnification is 2x (Upper panels) and 40x (lower panels). With age, undulation of the hepatic capsule and perivascular lymphoid aggregates (ii, iii -black arrows) are evident compared to 3-month old mice (i). Mild (v) and moderate (vi) increases in cell (anisocytosis) and nuclear size (anisokaryosis) were present in old individuals compared to 3-month old mice (iv). Additional histologic findings at 29 months (vi) included bile duct hyperplasia (white arrow), extramedullary hematopoiesis (black arrowheads), increased sinusoidal cellularity, and increased numbers of lipid-laden Ito cells. Single and double asterisks denote central veins and portal triads, respectively. (C) Properties of primary NSCs cultures with increasing animal age at the end of passages 1 and 3. (D-E) Multidimensional Scaling analysis results across datasets based on the breadth of the top 5% broadest H3K4me3 domains (D), or H3K27ac peak intensity at Super Enhancers only (E). (F-G) Principal Component Analysis results across datasets based on RNA expression (F), or H3K4me3 peak breadth (G).

**Figure S2: Separation of samples across tissues and cell types as a function of age (continued)**

Multidimensional Scaling analysis results across samples derived from specific tissues, Heart, Olfactory Bulb and primary NSCs cultures, based on RNA expression (A-C), H3K4me3 peak intensity (D-F), H3K4me3 peak breadth (G-I), H3K27ac peak intensity (all peaks: J-L; Super Enhancers only: M-O). Corresponding example of PCA analysis results across liver samples (P-T).

**Figure S3: Machine-learning analysis reveals that changes in enhancer score and H3K4me3 domain breadth with age can predict transcriptional aging (2-class classification)**

Scheme of the 2-class machine learning pipeline. Nnet: neural network, svm: support vector machine, rf: random forest, gbm: gradient boosting machine. (B-C) Classification accuracy over the 2-classes (*i.e.* upregulated vs. downregulated genes) cross tissues for Random Forest models (B) or Gradient Boosting models (C). The accuracy of the model trained in a specific tissue on the same tissue (*e.g.* the liver-trained model on liver data) is measured using held-out validation data, and for cross-tissue validation, the entire data of the other tissue was used. ‘Random’ accuracy is displayed to illustrate the accuracy of a meaningless model (∼50%). All tests were significantly more accurate than random. (D-E) Feature importance from Random Forest models (D; Gini and mean decrease in accuracy) or Gradient Boosting models (E; Gini). High values indicate important predictors. See analysis of 3-class models in **Figure 3**. Note that 2-class models systematically outperformed 3-class models, which is consistent with the increased complexity of a classification problem with the number of classes to discriminate.

**Figure S4: Age-related transcriptomic and epigenomic remodeling**

(A) Table of the number of genes or loci with significant age-remodeling across cells and tissues (FDR<5%). (B-D) UCSC Genome Browser Shots for example significantly remodeled loci in the liver samples. (E) Circular genome plot showing the genomic distribution of regions with significantly remodeled H3K4me3 breadth, H3K4me3 intensity or H3K27ac intensity in Cerebellum and Liver. Note that there is no obvious clustering on a specific chromosome. (F) Heatmap of genes with significant tissue-independent transcriptional changes with aging plotted in each of the tissues as called by DESeq2 with a significance threshold of FDR < 5%. Also see **Supplementary Table S4**. (G-I) Functional enrichments using the mHG test at for differential RNA expression (C), for differential H3K4me3 intensity (D), and for differential H3K27ac intensity (E). Note that none of the KEGG 2017 pathways were significantly enriched using gene with differential H3K4me3 breadth. Enriched pathways are plotted if 4 out of the 6 tests (RNA) or 3 out of the 5 tests (chromatin marks) were significant (FDR < 5%).

**Figure S5: CIBERSORT pipeline and signature matrix validation**

(A) Scheme of the CIBERSORT analysis pipeline. The signature matrix was built using publicly available RNA-seq datasets derived from pure cell types that are expected in the tissues under study (**Supplementary Table S2B**). (B) Heatmap of predicted cell type fractions on held-out purified RNA-seq samples using trained CIBERSORT Signature Matrix (see methods). (C) Heatmap of predicted cell type fractions on RNA-seq *in silico* mixtures of known cell types using trained CIBERSORT Signature Matrix (see methods).

(D) RNA deconvolution analysis of hippocampus RNA-seq datasets derived from in mouse models with known increased inflammatory cell content (in particular microglia) and activity. (Left panel) CK-p25 Alzheimer’s mouse model (Gjoneska et al. 2015); (Right panel) 5XFAD Alzheimer’s mouse model following forced 40Hz gamma oscillations (Iaccarino et al. 2016). Significance in one-sided Wilcoxon test. Also see **Supplementary Table S7C**. (E) RNA deconvolution analysis of tissue aging RNA-seq datasets. Significance in one-sided Wilcoxon test between 3 months and 29 months samples. Linear regression tests as a function of age were also non-significant. Also see **Supplementary Table S7D**. (F-G) Boxplots of DEseq2 normalized counts (log2 scale) for expression of immune marker genes *Ptrpc/CD45* and *Itgam/CD11b*.

**Figure S6: Conservation of age-related transcriptional deregulation in vertebrates**

(A) Functional enrichments using the mHG test at for differential RNA expression with aging in mouse, rat, human and African turquoise killifish samples. The mouse data is a subset of the panel shown in **Figure S4G** and is plotted as a comparison point. (B) Log2 Fold Change per unit of time during aging for genes orthologous to differentially expressed mouse genes in this study in rat and human (GTEX) samples, combining male and female samples. Significance in on sample one-sided Wilcoxon test the fold changes and 0 (*i.e.* no change with age). (C-D) Log2 Fold Change per unit of time during aging for genes orthologous to differentially expressed mouse genes in this study in human samples (GSE61260) only in males (C) or in combined male and female samples (D) after inclusion of metabolic parameters as covariates in limma (*i.e.* BMI and liver disease status). Significance in on sample one-sided Wilcoxon test the fold changes and 0 (*i.e.* no change with age).

**Supplementary Tables**

**Supplementary Table S1**: Sample design and biological information

**Supplementary Table S2**: Accession numbers of publicly available datasets used in this study **Supplementary Table S3**: Summary of machine-learning accuracy, specificity and sensitivity across algorithms

**Supplementary Table S4**: Genes with significant tissue-independent transcriptional changes with aging as called by DESeq2 with a significance threshold of FDR < 5%.

**Supplementary Table S5**: Functional annotation notes (IPA and KEGG)

**Supplementary Table S6**: HOMER analysis of repeat element transcriptional changes with aging.

**Supplementary Table S7**: Results CIBERSORT RNA deconvolution algorithm.

